# Loss of Tet2 affects proliferation and drug sensitivity through altered dynamics of cell-state transitions

**DOI:** 10.1101/2020.01.20.911834

**Authors:** Leanna Morinishi, Karl Kochanowski, Ross L. Levine, Lani F. Wu, Steven J. Altschuler

## Abstract

A persistent puzzle in cancer biology is how mutations, which neither alter canonical growth signaling pathways nor directly interfere with drug mechanism, can still recur and persist in tumors. One notable example is the loss-of-function mutation of the DNA demethylase Tet2 in acute myeloid leukemias (AMLs) that frequently persists from diagnosis through remission and relapse (Rothenberg-Thurley *et al.*, 2018; Corces-Zimmerman *et al.*, 2014; Nibourel *et al.*, 2010), but whose fitness advantage in the setting of anti-leukemic chemotherapy is unclear. Here we use paired isogenic human AML cell lines to show that Tet2 loss-of-function alters the dynamics of transitions between differentiated and stem-like states. Mathematical modeling and experimental validation reveal that these altered cell-state dynamics can benefit the cell population by slowing population decay during drug treatment and lowering the number of survivor cells needed to re-establish the initial population. These studies shed light on the functional and phenotypic effects of a Tet2 loss-of-function in AML, illustrate how a single gene mutation can alter a cells’ phenotypic plasticity, and open up new avenues in the development of strategies to combat AML relapse.

## Introduction

A major challenge in cancer biology is to understand the function of recurrent mutations in the emergence of tumors or response to drug therapy. Some mutations clearly benefit cancer populations either by 1) altering regulation of growth signaling or programmed cell death (e.g. mutations in p53 or TGFβ signaling, Sanchez-Vega, *et al.*, 2018) or by 2) directly interfering with drug effect (e.g. acquired EGFR T790M resistance mutations in response to EGFR tyrosine kinase inhibitor therapy, Ma, Wei and Song, 2011). However, for many observed mutations it is unclear how they affect either proliferation or drug resistance.

One example are mutations in the DNA demethylase Tet2, found in ~15-20% of *de novo* AML patients (Nibourel *et al.*, 2010; Metzeler *et al.*, 2011; Cancer Genome Atlas Research Network *et al.*, 2013, Moran-Crusio *et al.*, 2011). Mutated Tet2 is associated with mutational persistence and adverse outcome in human AML (Rothenberg-Thurley *et al.*, 2018; Ding *et al.*, 2011). However, due to its ability to alter DNA methylation genome-wide, the mechanisms by which Tet2 mutation confers a benefit to AML cancer cell populations remain unclear. Here we investigate this puzzle using an integrated approach of mathematical modeling and experimentation in paired WT and Tet2-mutant isogenic human AML cell lines. We discover that Tet2 mutation alters the dynamics of transitions between distinct stem-like and differentiated cell states, which enhances population fitness in chemotherapy and lowers the number of cells needed to establish a cell population.

## Results

### Tet2 loss-of-function mutation renders AML cell populations more stem-cell like

To investigate the consequences of Tet2 mutation, we chose to compare two pairs of isogenic human myeloblast cell lines, each expressing wildtype or mutant Tet2 (Figure 1A). The AML cell lines KG1 and Thp1 (Kunimoto *et al.*, 2018) were selected as they express wildtype Tet2 (Tet2^WT^) but do not express mutant FLT3, which is known to have synergistic epigenetic effects (Shish *et al.*, 2015). In human AMLs, Tet2 is often mutated in AML with truncating mutations or missense mutations in its catalytic domain (Hirsch *et al.*, 2018), resulting in Tet2 loss-of-function (Smith *et al.*, 2010). Therefore, isogenic cell lines were created by knocking out Tet2 in the chosen cell lines (Tet2^KO^, Methods). Loss of wildtype Tet2 expression was confirmed *via* RT-qPCR of the Tet2 transcript and immunoblotting of the N-terminus of the protein (Supplemental Figure 1). To confirm that loss of Tet2 has the expected effect on DNA methylation (Yamazaki *et al.*, 2015; Asmar *et al.*, 2013; Rasmussen *et al.*, 2015), we performed DNA methylation profiling. As expected, Tet2^KO^ cell lines display a significantly higher degree of overall hypermethylation (t-test, p-values: KG1 <2.2e-16 and Thp1 2.9e-4) compared to their WT counterparts, with high reproducibility across replicates (Figure 1B, Supplemental Figures 2A, 3).

**Figure 1.**
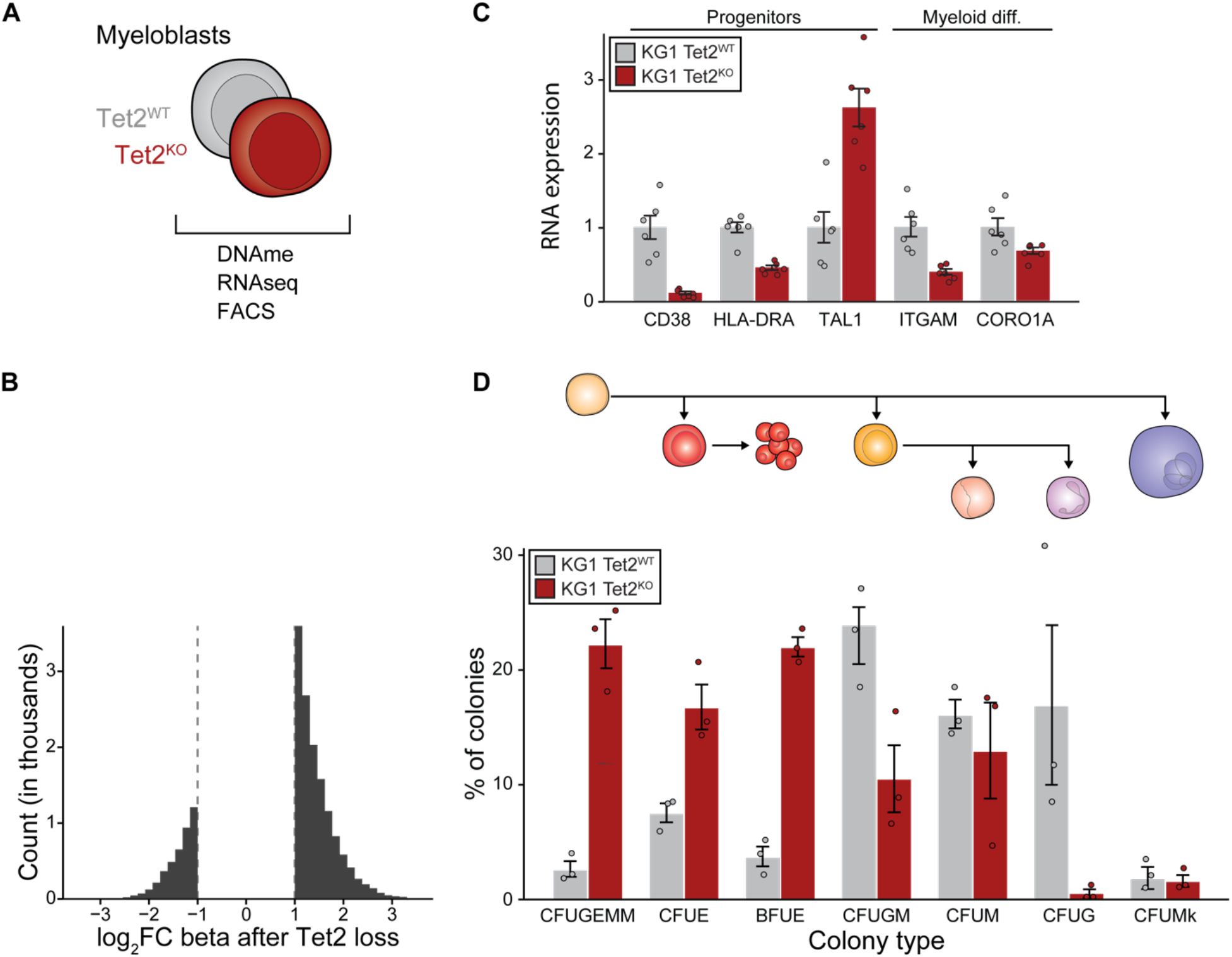
Tet2^KO^ cells are more stem-like than Tet2^WT^ isogenic counterparts. (A) Overview of molecular profiling performed on isogenic Tet2 mutant AML cell lines. (B) Tet2^KO^ cell lines are more hypermethylated than their wildtype counterparts. Shown is a histogram of log_2_fold-change (CpG beta values) in Tet2^KO^ cell lines for differences greater than 2-fold from a paired analysis. (C) Genes that modulate differentiation and stemness are differentially expressed as measured by RNAseq after Tet2^KO^ in both KG1 and Thp1 (Supplemental Figure 2C, mean ± s.e., n=6). Tet2^KO^ cell lines show decreased expression of myeloid commitment markers (ITGAM, CORO1A) and increased expression of markers associated with stemness (CD38^-^, HLA-DRA^-^, TAL1) compared to Tet2^WT^. (D) KG1 and Thp1 (Supplemental Figure 4A) Tet2^KO^ cell lines produce a higher percentage of colonies associated with oligopotent progenitor cells (CFUGEMM) compared to Tet2^WT^ cells (mean *±* s.e., n=3) in methylcellulose colony forming assays.

To gain an unbiased overview of the molecular changes induced by Tet2^KO^, we examined the transcriptomic and epigenomic profiles of Tet2 WT and KO cell populations. RNAseq analysis identified ~300 gene transcripts that are similarly differentially expressed in both Tet2^KO^ cell lines (fold-change>2, Supplemental Figure 2B). Notably, reduced expression of myeloid differentiation markers (ITGAM, CORO1A, Wald test, BH adjusted p-values 9.99e-9 and 7.73e-2, Figure 1C, Supplemental Figure 2C) and increased expression of markers associated with leukemic progenitor cells (CD38^-^, HLA-DRA^-^, TAL1, Wald test, BH adjusted p-values 3.83e-26, 1.74e-16, and 3.90e-7, Figure 1C, Supplemental Figure 2C) (Nishioka *et al.*, 2013) were observed. These data are consistent with analysis of TCGA LAML and OHSU datasets, which showed that Tet2 expression is strongly correlated with genes expressed in the granulocytic lineage (Fisher’s exact test, adjusted p-values 8e-25 and 3.7e-19, Supplemental Tables 1–2, Methods).

Differentially expressed genes in Tet2^KO^ cells were found to be highly enriched for targets of Runx1, a hematopoietic regulator known to promote stemness and myeloid fate decisions (Fisher’s exact test, p-value 7.76e-6, Supplemental Figure 2D, Supplemental Tables 3–4) (Kuleshov *et al.*, 2016; Ran *et al.*, 2013). Consistently, analysis of differentially methylated regions showed the proximal promoter of Runx1 to be significantly affected in Tet2^KO^ (Wald test, minimum adjusted p-value 1.22e-27, Supplemental Figure 2E), with a concomitant increase in the expression the Runx1a isoform (Supplemental Figure 2F). Methylcellulose colony forming unit assays of Tet2^KO^ cells also show increased numbers of colonies associated with oligopotent progenitor cells compared to Tet2^WT^ controls (CFUGEMMs, Figure 1D, Supplemental Figure 4A). Overall, these data suggested that Tet2^KO^ populations acquire more stem-like signatures, which was further validated by comparing their DNA methylation profiles to those of normal hematopoietic progenitors and leukemic stem cells (LSCs, Supplemental Figure 4B,C) (Horvath, 2013; Jung *et al.*, 2015).

The Tet2^KO^-mediated acquisition of stem-like signatures was much more pronounced in KG1 than Thp1 in terms of expression profile and potential to form diverse myeloid lineages in colony-forming assays (Supplemental Figure 4). This is presumably because the monocyte-like cells, Thp1, are already more differentiated than the myeloblast-like cells, KG1, regardless of Tet2 mutational status. We therefore chose to focus on KG1 cells for the remainder of this work.

### Tet2 mutation changes dynamics of cell-state switching

To test whether this change in stem-like molecular signatures is caused by an increase in the fraction of cells with CD34^hi^CD38^lo^ surface marker expression (a classic LSC-like profile (Nishioka *et al.*, 2013)) in Tet2^KO^ compared to Tet2^WT^ cells, we quantified CD34 and CD38 signals in the population using flow cytometry. Quantification of KG1 CD34/CD38 expression revealed that the fraction of CD34^hi^CD38^lo^ cells was indeed increased in Tet2^KO^ cell populations (Figure 2A).

**Figure 2.**
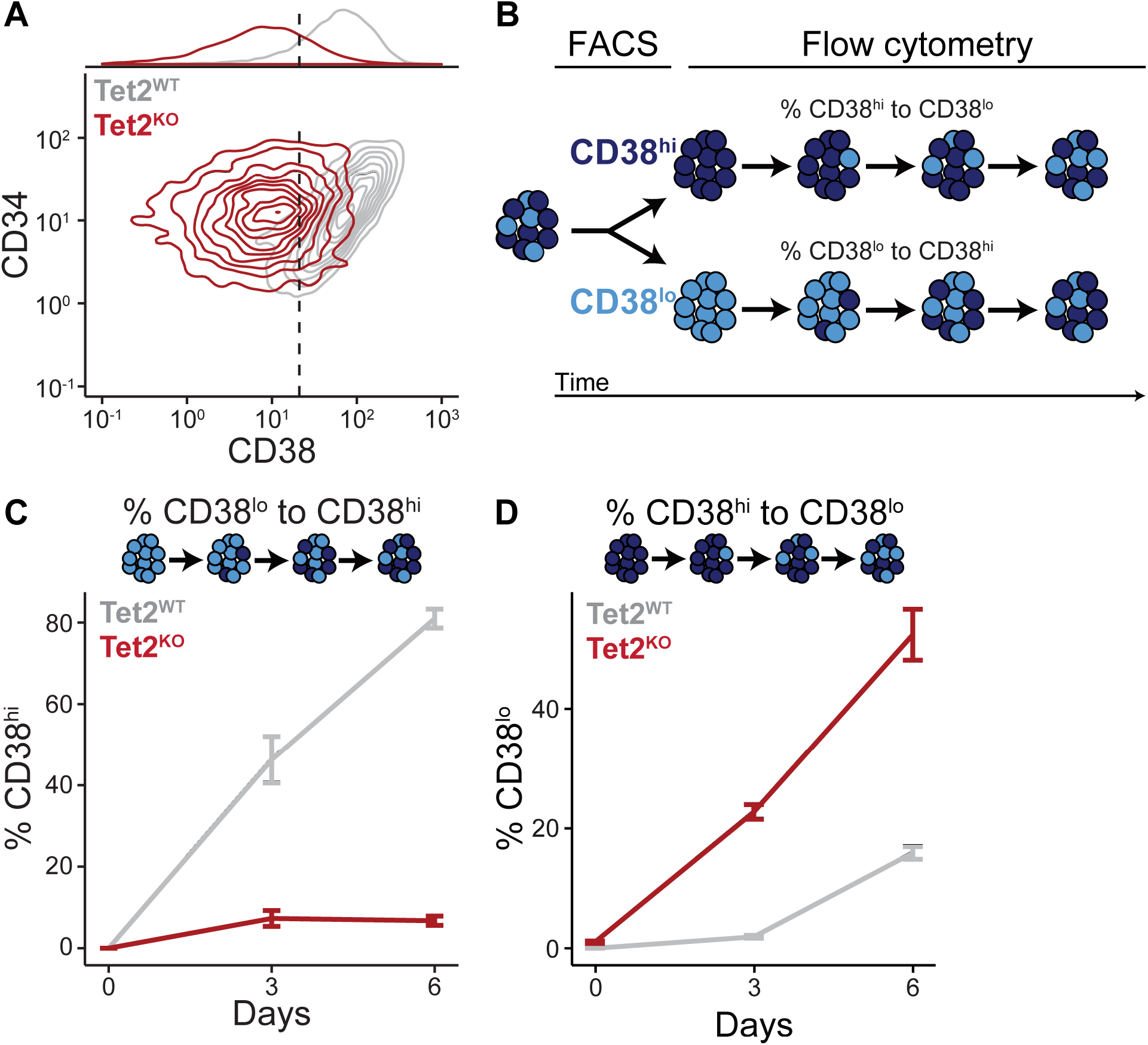
Tet2^KO^ increases the propensity of differentiated (CD34^hi^CD38^hi^) Cells to switch to a more stem-like (CD34^hi^CD38^lo^) cell state. (A) KG1 Tet2^KO^ cell lines show a shift in CD38 surface marker expression. (B) Schematic illustration of phenotypic segregation and flow cytometry experiment to quantify CD38 expression over time. (C-D) The percent CD38^hi^ cells in an originally pure population of sorted CD38^lo^ cells (C) or the percent CD38^lo^ cells in an originally pure population of sorted CD38^hi^ cells (D) after 0, 3, and 6 days in untreated conditions (mean ± s.e., n=3).

How does Tet2^KO^ alter the ratio between stem-like and differentiated cells? One possibility is that Tet2^KO^ simply reduces the rate of differentiation. To test this, subpopulations of stem-like (*L*, CD34^hi^CD38^lo^) and differentiated (*H*, CD34^hi^CD38^hi^) cells were sorted in both Tet2^WT^ and Tet2^KO^ populations, and the ratio of *L* and *H* cells was monitored over time (Figure 2B, Supplemental Figure 5A). Indeed, sorted stem-like cell populations repopulated the differentiated state more slowly in Tet2^KO^ (Figure 2C, Supplemental Figure 5B). Surprisingly, in Tet2^KO^, the sorted differentiated cell populations repopulated the stem-cell like state more rapidly than Tet2^WT^ populations (Figure 2D, Supplemental Figure 5B). These data suggest that Tet2^KO^ facilitates the reversible switching from differentiated to stem-like states in AML cells and provides intuition for why Tet2^KO^ has a larger fraction of stem-like cells in the population.

### Mathematical modeling illustrates consequences of altered cell-state dynamics for survival of cell population

What are the functional consequences of having such altered cell-state dynamics, and in particular a higher fraction of stem-like cells? To address this question, we developed a general mathematical model with three cell states (Figure 3, second panel): a stem-cell-like state *L,* a differentiated state *H,* and an irreversible cell death state (to account for the impact of drug treatment). This model is fully characterized by a linear, homogeneous system of ordinary differential equations (ODEs) with 6 parameters with values ≥ 0 (Methods). Parameters are denoted by: *d_L_* and *d_H_* the death rates of the *L* and *H* states, respectively; *g_L_* and *g_H_*, the rates at which the *L* and *H* states proliferate; and *r_LH_* and *r_HL_* the transition rates from *L* to *H* and *H* to *L*, respectively.

**Figure 3.**
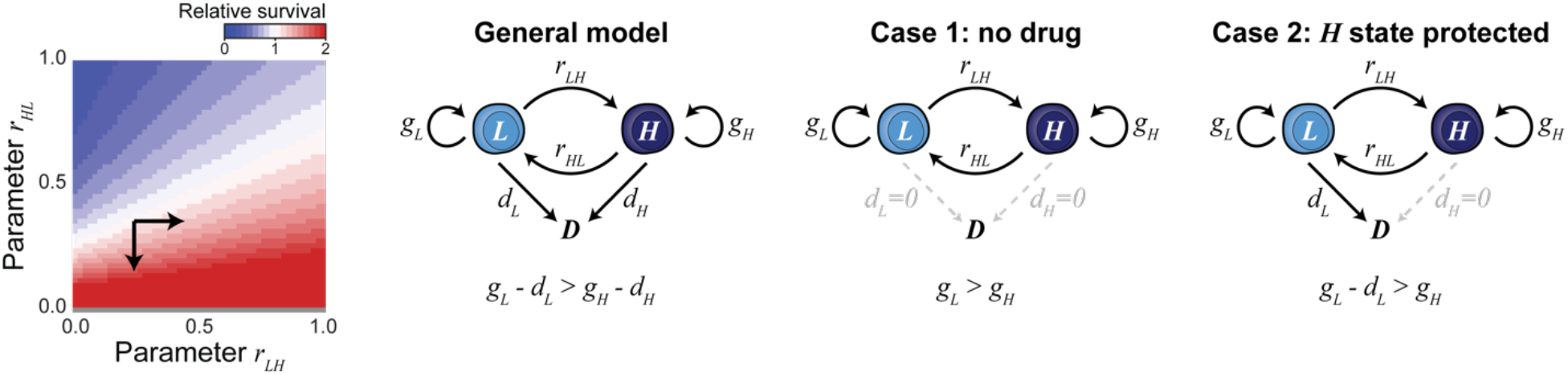
Mathematical model reveals advantageous and disadvantageous parameter regimes for cell-state switching. (left) If model parameters satisfy Inequality 1, both increasing ***r_HL_*** and/or decreasing ***r_LH_*** (black arrows) slows population decay and benefits a drug-treated cancer population. (General model) Schematic representation of mathematical model with three cell states: a stem-cell-like state ***L***, a differentiated state ***H***, and an irreversible cell death state. Parameters ***d_L_*** and ***d_H_*** represent the death rates of the ***L*** and ***H*** states; ***g_L_*** and ***g_H_*** the proliferation rates; and ***r_LH_*** and ***r_HL_*** the transition rates from ***L*** to ***H*** and ***H*** to ***L***. (Case 1) If cells rarely die (**d_L_** ≈ 0, **d_H_** ≈ 0) and the proliferation rate ***g_L_*** is higher than ***g_H_***, increasing ***r_HL_*** is always beneficial. (Case 2) Conversely, if one state were protected from drug effect (**d_H_** ≈ 0), increasing ***r_HL_*** is only beneficial in the unlikely scenario when net production rate of the other state ***L*** is larger than its proliferation rate ***g_H_***.

Using this model, we asked under which circumstances the observed altered cell-state dynamics – specifically an increased switch rate towards a stem-like state – would benefit a cell population. First, we focused on the impact of such altered cell-state dynamics during drug treatment (i.e. high death rates). In this case, the time to population collapse is dominated by the largest negative eigenvalue; the less negative the eigenvalue, the slower the population collapse. By computing the eigenvalues for the ODE, it can be seen that either increasing *r_HL_* and/or decreasing *r_LH_* slows population decay and “benefits” a drug-treated cancer population as long as (Figure 3, first panel; Methods):

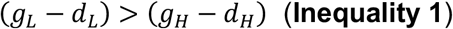

This inequality simply compares net production rates (proliferation rate minus death rate) and requires it to be higher at *L* than *H.* In fact, as long as Inequality 1 is true, increasing *r_HL_*, or decreasing *r_LH_* will eventually slow population decay (arrows in phase plane diagram below; see Methods for details). Thus, as long as the net production rate of the stem-like state L exceeds that of the differentiated state H, altered cell state dynamics (i.e. towards a higher fraction of stemlike cells) will always benefit the cell population. This finding is in line with recent studies suggesting the possibility that cancer cells can reversibly transit between stem and differentiated states with different drug sensitivities (Jordan *et al.*, 2016; Su *et al.*, 2017; Gupta *et al.*, 2011).

Importantly, the same inequality also captures other cases in which switching to a stemcell-like state is either beneficial or detrimental (Figure 3, third and fourth panels). If cells rarely die, such as in a non-drug-treated condition (*d_L_* ≈ 0, *d_H_* ≈ 0) (**Case 1**), inequality 1 simplifies to

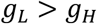

Here, switching to the *L* state (increasing *r_HL_*) is beneficial for the cell population when the proliferation rate *g_L_* of *L* is higher than the proliferation rate *g_H_* of *H*, which is likely given the lower propensity of differentiated cells to divide. In this case, we expect cells carrying mutations that increase *r_HL_* to take over the population. Conversely, if the differentiated cell state *H* were protected from the adverse effects of drug treatment (*d_H_* ≈ 0) (**Case 2**), inequality 1 simplifies to

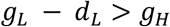

Here, increasing *r_HL_* is only beneficial for the cell population if the net production rate of *L* is larger than the proliferation rate *g_H_* of the differentiated state *H*, which is unlikely for high doses of drug (*d_L_* » 0). In this case, we expect cells carrying mutations that increase *r_HL_* to be depleted from the population over time.

Overall, this general mathematical model shows that altered cell-state dynamics – specifically an increased switch rate towards a stem-like state – can indeed benefit the cell population (both in presence or absence of drug treatment), as long as it falls within a parameter regime outlined by inequality 1 (i.e. net production rate of stem-like state exceeds that of the differentiated state).

### Experimental validation of modeling predictions

Does the Tet2 mutation put the AML cancer population in this predicted advantageous parameter regime? Since the proliferation, death and switching rates cannot be disentangled directly from measurements, we estimated these rates by fitting our model to time course data of sorted Tet2^KO^ and WT populations in the presence of cytosine arabinoside (AraC, a common first-line chemotherapy drug for AML) (see Methods, Figure 4B). These estimates confirmed that the proliferation rate during drug treatment of stem-like cells far exceeds that of differentiated cells, both for Tet2^KO^ and WT populations (Figure 4A-B). Thus, both populations are in a regime where the net production of stem-like cells outstrips the net production of differentiated cells in drug treatment (as required by Inequality 1), and therefore switching to the stem-like state is more advantageous. The Tet2^KO^ populations have a higher *H → L* switching rate and a lower *L → H* switching rate; hence, the model predicts that the Tet2^KO^ population, as a whole, is more drug resistant. Indeed, drug treatment experiments confirmed that Tet2^KO^ populations are less sensitive to AraC and doxorubicin than Tet2^WT^ populations (Figure 4C, Supplemental Figure 5C).

**Figure 4.**
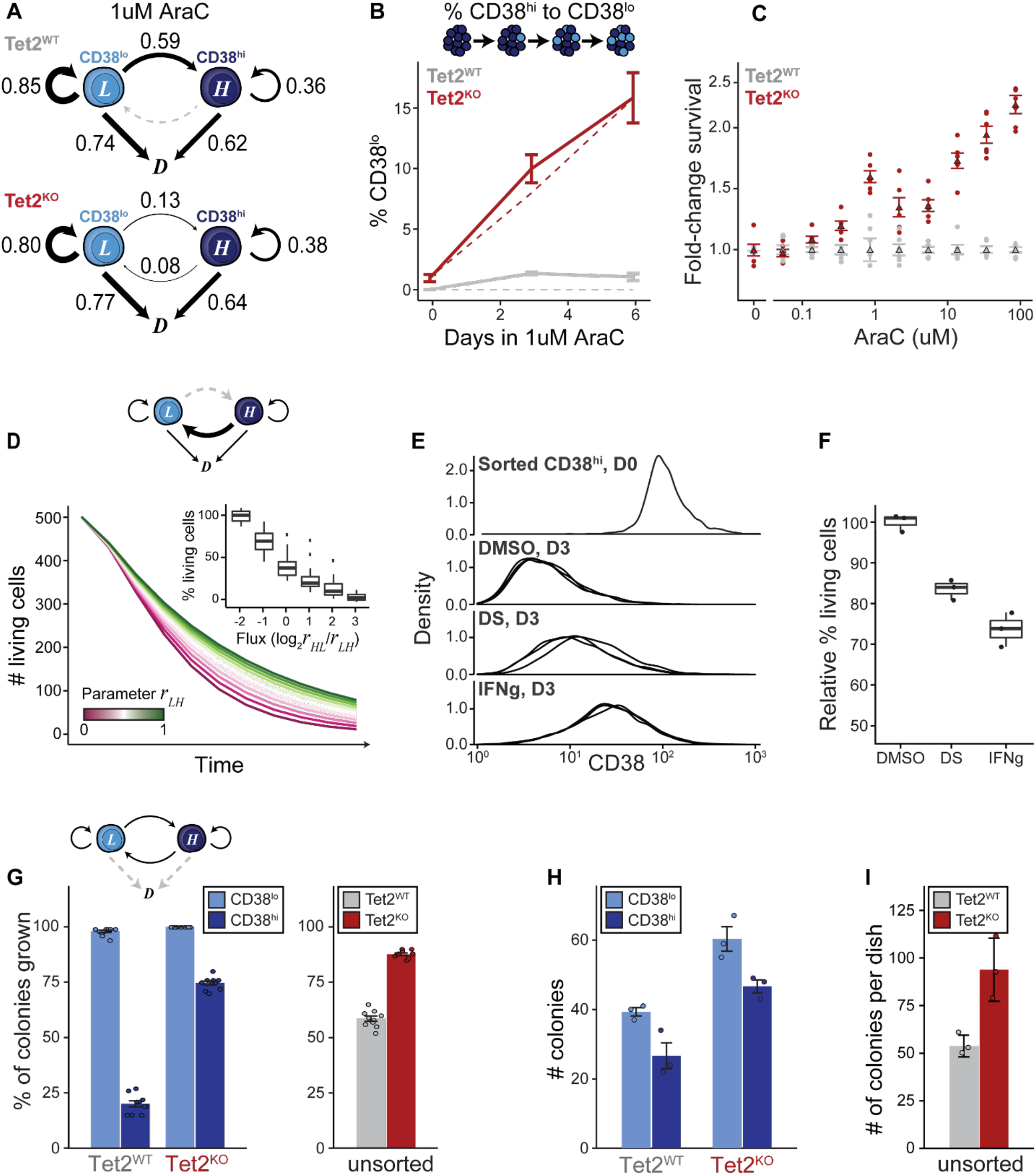
Tet2^KO^ cells’ altered cell-state dynamics enable longer-term drug survival in chemotherapy. (A) Schematic illustration of model of cell-state transition, cell proliferation, and cell death in 1uM AraC treatment. Numbers denote corresponding estimated rates of transition, proliferation, and death. (B) The percent of CD38^lo^ cells in an originally pure population of sorted CD38^hi^ cells in 1uM AraC after 0, 3, and 6 days (mean ± s.e., n=3). Dashed lines represent predictions from the model. (C) The viability of cells treated with 72h of varying concentrations of AraC were assessed by CellTitreGlo. Shown is fold-change viability relative to Tet2^WT^ cells as a function of chemotherapy concentration (mean ± s.e., n=5). (D) Model predictions for cell drug survival given increasing values of the transition parameter *r_HL_* for a toy model that fulfills Inequality 1. Plotting survival as a function of the ratio of *r_HL_/r_LH_* (inset). (E) CD38^hi^ cells were isolated by FACS, then incubated with or without effector for 3 days in the presence of 1 uM AraC. Shown are distributions of sorted Tet2^KO^ CD38^hi^ cells after 3 days in DMSO, DS, or IFNg treatment (n=3, Tet2^WT^ in Supplemental Figure 6A). (F) Sorted CD38^hi^ cells were subjected to 1uM AraC treatment with or without effector. The percent of living cells in the population relative to control after 3 days is shown on the y-axis (mean ± s.e., n=3). (G) Model predictions for colony outgrowth from a small population given sorted CD38^lo^ and CD38^hi^ or unsorted cells (see Methods, mean ± s.e., 10 runs). (H) Number of colonies in a methylcellulose colony forming unit assay for Tet2^WT^ and Tet2^KO^ cells after seeding equal numbers and 14d incubation (mean ± s.e., n=3). (I) Number of colonies in methylcellulose assay Tet2^WT^ and Tet2^KO^ cells sorted for CD38 expression (mean ± s.e., n=3).

Further analysis of the model suggested two additional predictions. First, the model suggested that a population with a higher fraction of differentiated cell states will show reduced population survival in drug treatment (Figure 4D). To test this conjecture, two effectors known to enrich AML cell populations for the differentiated state (without altering cell death, Supplemental Figure 6C) were used, namely the inflammatory stimulus interferon gamma (IFNg) and the aldehyde dehydrogenase inhibitor disulfiram (DS) (Amici *et al.*, 2018; Xu *et al.*, 2017). We confirmed that treatment with IFNg or DS enriched both Tet2^KO^ and WT populations for the differentiated cell state after 72 hours of exposure (Methods, Figure 4E, Supplemental Figure 6A-B). Moreover, both effectors increased the efficacy of AraC treatment in both Tet2^KO^ and WT populations, while several effectors not known to affect cell-state transitions did not alter AraC sensitivity (Figure 4F, Supplemental Figure 6C, Supplemental Table 5).

Second, the model predicted that the increased ability of Tet2^KO^ to revert to a stem-like cell state (with its higher proliferation potential), will allow a Tet2^KO^ population to better regrow out of drug than a Tet2^WT^ population (Figure 4G). To test this prediction, we compared the number of colonies formed by Tet2^KO^ and Tet2^WT^ populations in methylcellulose assays, which assesses the ability of isolated cells to reform a colony (Figure 4H). As predicted, sorted stem-like cell populations showed increased cell colony numbers in both Tet2^KO^ and Tet2^WT^ populations. Moreover, unsorted Tet2^KO^ populations formed approximately 2x more colonies than their Tet2 WT counterparts (Figure 4I), highlighting their increased potential for population renewal. Thus, our results suggest that Tet2^KO^ populations have a higher likelihood to regrow an AML population from few surviving cells in the setting of therapeutic perturbations.

## Discussion

Taken together, our results reveal that mutation of the epigenetic modifier Tet2 in AML cell populations alters the transition dynamics between stem-like and differentiated cell states. These altered cell-state dynamics confer several benefits to the population. First, Tet2 mutation enables differentiated cells to switch to a stem-like state whose increased drug sensitivity is compensated by a higher proliferation rate, thus increasing population survival. Second, Tet2 mutation increases the propensity to regrow an AML population after drug treatment from few surviving cells, due to the increased ability of Tet2^KO^ cells to revert back to a proliferative stemlike cell state (Figure 4A). These findings provide a rationale for why the detection of Tet2 mutants in patients in remission is strongly associated with a higher chance of relapse (Rothenberg-Thurley *et al.*, 2018; Ding *et al.*, 2012).

This work has important implications for AML mutations beyond Tet2: the mathematical model shows that as long as stem-like and differentiated cell states differ in their net cell production during drug treatment (i.e., Inequality 1 is fulfilled), any other mutation increasing the transition rate towards the stem-like state will be beneficial as well. Thus, we conjecture that mutations in other epigenetic modifiers might confer a fitness advantage to drug-treated AML populations through similar mechanisms.

More generally, this work provides further evidence that mutations, which modulate switching rates between more and less drug-resistant states, may provide an “evolutionary shortcut” to counteract the adverse effect of drug treatment (Jordan *et al.*, 2016; Su *et al.*, 2017; Gupta *et al.*, 2011). In cancer, the target space for mutations that directly interfere with drug function is likely smaller than the target space for dysregulation of cell states; this is known to be the case in bacteria, where the number of genes conferring increased antibiotic tolerance far exceeds the number of genes conferring bona fide drug resistance (Girgis, Harris, and Tavazoie, 2012; Brauner *et al.*, 2016). Future studies into identifying cell states and dynamics of transitions between these states will provide new insight into why certain mutations are selected for, and persist in, cancer, and led to new therapeutic approaches which increase the efficacy of current cancer therapies.

## Acknowledgments

Thanks to Kevin Shannon, Dylan Altschuler, Jason Altschuler, Jake Bieber, Jeremy Chang, Janice Goh, Louise Heinrich, Weiyue Ji, Capria Rinaldi, and Xiaoxiao Sun for helpful discussions and feedback. Funding sources: L.M. is supported by NSF GRFP; K.K. is supported by the European Molecular Biology Organization (long-term fellowship ALTF 1167–2016) and the Swiss National Science Foundation; R.L.L. is supported by National Cancer Institute grants P01 CA108671 and R35197594 as well as a Leukemia & Lymphoma Society Specialized Center of Research grant; L.F.W. is supported by NCI-NIH RO1 CA185404 & CA184984; and S.J.A. is supported by GM112690, NSF PHY-1545915 and SU2C/MSKCC 2015-003.

## Author contributions

Conceptualization and Methodology, L.M., K.K., S.J.A. and L.F.W.; Investigation, Formal Analysis, and Visualization, L.M.; Resources, R.L.L.; Writing – Original Draft, L.M. and K.K.; Writing – Review & Editing, L.M., K.K., S.J.A. and L.F.W.; Supervision, K.K., R.L.L., S.J.A. and L.F.W.

## Declaration of interests

R.L.L. is on the supervisory board of Qiagen and is a scientific advisor to Loxo, Imago, C4 Therapeutics and Isoplexis which include equity interest. He receives research support from and consulted for Celgene and Roche and has consulted for Lilly, Janssen, Astellas, Morphosys and Novartis. He has received honoraria from Roche, Lilly and Amgen for invited lectures and from Gilead for grant reviews.

## Materials & Correspondence

Correspondence to Steven J. Altschuler and Lani F. Wu.

## STAR Methods

### Cell-state transition model

We modeled a cell system with ***m*** states as a linear, homogenous system of ODEs. In matrix form this is written as

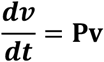

where ***v*** is cell-state vector of length ***m***, and ***P*** is a ***m x m*** matrix representing cell-state transitions. We do not assume the matrix ***P*** is stochastic, which allows expansion or contraction of the total number of cells in the system—reflecting cell division or drug sensitivity—to occur. In our study, we considered three states: *L* (CD38^lo^), ***H*** (CD38^hi^), and ***D*** (dead):

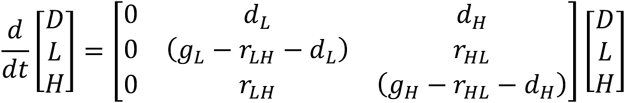

where ***g_L_*** and ***g_H_*** are the growth rates and ***d_L_*** and ***d_H_*** the death rates of cell-states ***L*** and ***H***, respectively. Here, ***r_LH_*** is the transition rate from ***L*** to ***H***, and ***r_HL_*** is the transition rate from ***H*** to ***L***. All matrix parameters are considered to be real, non-negative numbers. If we define

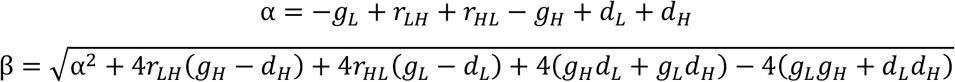

then the eigenvalues of *P* are

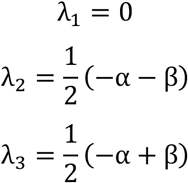

with eigenvectors

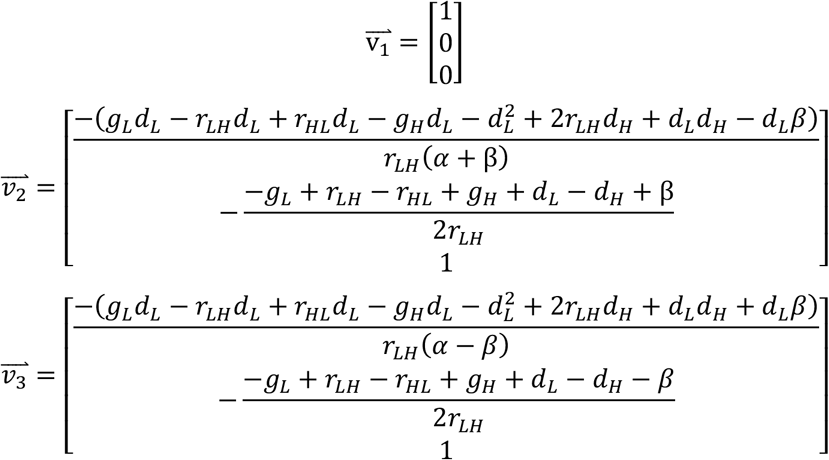

The first eigenvalue corresponds to a “trivial” solution in which dead cells remain dead. For convenience, we define the difference in production between the two states

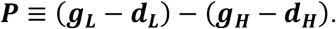

To find the regime where changing *r_HL_* (asympototically) increases survival rate, we focus on the larger of the other two eigenvalues, namely **λ_3_**. Taking the derivative of **λ_3_** with respect to the state-switching parameters gives:

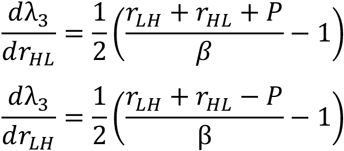

Using Wolfram|Alpha, we find: a) 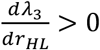 holds if *P* > 0 and 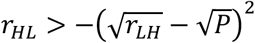; and b) 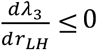 holds if either (i) *P* > 0, 0 ≤ *r_LH_* < *P*, and 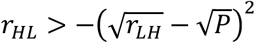, or if (ii) *P* > 0 and *r_LH_* ≥ *P*. In our model, we only consider non-negative, real-valued parameter values. Thus, as long as the inequality *P* > 0 holds, 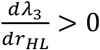 and 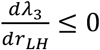.

To find best fit parameters for this system, cell-state compositions from the flow cytometry experiment (described above) were used as input into the model. Prediction error at each time step was minimized with fminsearch in MATLAB (version R2019a) using data from both CD38^hi^- and CD38^lo^-sorted subpopulations. Solving the matrix ODE gave

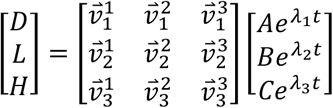

After solving for the constants ***A, B***, and ***C***, model predictions were made using the best fit parameters and selected initial conditions, unless otherwise noted.

For colony forming simulations, 100 individual cells were “seeded” as colony founders with 50% in the ***L*** state and 50% in ***H*** unless otherwise noted. To simulate growth without drug treatment, rates were estimated by fitting the model to flow cytometry experimental data (described above) without AraC treatment. The number of cells in the ***L, H***, or ***D*** states at time ***t*** was found with solution to the matrix ODE at time ***t***. Remaining fractions of cells were treated as the probability of an additional cell transition. Cells “grew” for 10 time-steps before the number of cells in each colony was counted. Colonies with more than 50 cells were considered “grown out”.

### Cell lines, reagents, and cell culture

KG1 cells and Thp1/Thp1 Tet2^KD^ cells were generous gifts from the Shannon and Levine labs respectively. Cells were cultured in ATCC-recommended media and incubated at 37°C with 5% CO_2_. All cell lines were maintained in 75mL or 250mL Suspension Culture flasks (CellTreat, Pepperell, MA) between 0.5e6 and 1e6 cells/ml.

To make Tet2^KO^ cells in the KG1 cell line, guides targeting exon 3 of Tet2 (an exon with frequent indel mutations in patients with myelodysplastic syndrome (MDS) (Smith *et al.*, 2010)) were designed in Benchling (5’-CACCGAGGCCAATTAAGGTGGAACC-3’). pSpCas9(BB)-2A-Puro (PX459) V2.0 was a gift from Feng Zhang (Addgene plasmid #62988). Guides were cloned into PX459 using BbsI sites, and vectors transfected into KG1 cells with the Cell Line Nucleofector Kit T (Lonza Group, Basel, Switzerland) with protocol T-020 and according to the manufacturer’s protocol.

Prior to addition to cell culture media, Cytosine Beta-D-Arabinofuranoside (Sigma-Aldrich, St. Louis, MO) and 2-Deoxy-D-glucose (Sigma-Aldrich) were dissolved in H2O, CCCP (Sigma-Aldrich), Disulfiram (Sigma-Aldrich), Etacrynic acid (Sigma-Aldrich), and Torin1 (Cell Signaling Technology, Danvers, Massachusetts) were dissolved in DMSO, and IFNg was dissolved in culture media to 1000x the desired final concentration. Final DMSO (or other diluent) concentration was always 0.1%.

### DNA methylation profiling

DNA methylation was profiled using Illumina’s Infinium MethylationEPIC BeadChip (Illumina, San Diego, CA). DNA of technical replicates for each condition was extracted using the Zymo Quick-DNA kit (Zymo Research, Irvine, CA, KG1 n=6 per condition, Thp1 n=2 per condition). Bisulfite conversion, nanodrop quantitation, array scanning, and normalization was performed by the Vincent J. Coates Genomics Sequencing Laboratory at UC Berkeley. Differential methylation analysis was performed with ChAMP (package version 2.14.0) in R (version 3.6.0). Differential methylated region analysis was performed with DMRcate (version 1.20), and visualization done with Gviz (version 1.28).

### mRNA-seq

RNA extraction of technical replicates was performed using the Lexogen SPLIT RNA extraction kit (Lexogen, Vienna, Austria, n=6 per condition), and libraries were prepared using the QuantSeq 3’ mRNA-Seq Library Prep Kit FWD for Illumina. Samples were quantified with Invitrogen Qubit (Invitrogen, Carlsbad, CA) prior to pooling, and library size and integrity was confirmed using the Agilent Bioanalyzer with the high-sensitivity DNA kit (Agilent, Santa Clara, CA). RNA sequencing was performed using 50bp single-end sequencing on the Illumina HiSeq 4000 in the Center for Advanced Technology at UC San Francisco. A PhiX control library was used as an in-run control, spiked in at 5%. Reads were mapped and counted using the Integrated QuantSeq data analysis pipeline on Bluebee Platform (Bluebee, Rijswijk, Netherlands). Briefly, reads were trimmed with BBDuk, aligned to human GRCh38 with STAR, and counted with HTSeq-count.

Gene filtering and differential expression analysis was performed in R. Genes were filtered by count such that all genes had 3 or more samples with 10 or more counts. Differential expression analysis was then performed using DESeq2 (version 1.24) on gene counts. Genes that were found to be significantly differentially expressed in a paired analysis of untreated cell lines were submitted to Enrichr (Kuleshov *et al.*, 2016) for enrichment analysis.

### Tet2 expression analysis

Expression profiles were queried from the Cancer Genomics Data Server (CGDS) using cdgsr (version 1.3.0). For each dataset (TCGA-LAML and AML-OHSU), genes with coefficient of variation < 0.1 were dropped from subsequent analysis. Pearson correlation calculations were performed in R. Genes with expression that strongly correlated (cor > 0.45) or anti-correlated (cor < −0.45) with Tet2 expression were submitted to Enrichr (Kuleshov *et al.*, 2016) for enrichment analysis.

### RT-qPCR and PCR

RNA was extracted from three technical replicates with the Direct-zol RNA Miniprep kit (Zymo Research) according to manufacturer’s instructions with TRI Reagent (Thermo Fisher Scientific, Waltham, MA). Reverse transcription with iScript Reverse Transcription Supermix (Bio-Rad, Hercules, CA) was followed by qPCR with DyNAmo Flash SYBR Green qPCR kit (Thermo Fisher Scientific) according to the recommended protocol. Primers were obtained from IDT (San Jose, CA). Primer sequences for human Runx1, Tet2 and GAPDH were as previously described^28^.

### Methylation age and Stemness score

Quantile normalized values from the aforementioned ChAMP analysis were used as input to the DNA methylation age calculator in R as described in the tutorial (horvath.genetics.ucla.edu) (Horvath, 2013). For visualization purposes, results for Tet2^WT^ and Tet2^KO^ cells were normalized to Tet2^WT^.

For the “stem-like” epigenetic signatures, we used a publicly available dataset of DNA methylation profiles from normal hematopoietic progenitors and leukemic cells sorted by CD34/CD38 expression (Jung *et al.*, 2015). Linear discriminant analysis (LDA) was applied using the MASS package (version 7.3-51.4) in R to identify an optimal transform that increased separation of DNA methylation profiles across three key reference populations: normal HSCs, CD34^+^CD38^-^ putative LSCs, and CD34^-^ leukemia cells. The rest of the public dataset, as well as methylation data from the paired Tet2^WT^/Tet2^KO^ cell lines, was projected into the lower-dimensional LDA space. The final “stemness score” was simply the LD2 value of the query methylation profile in LD space. A constant was added to the final score for visualization purposes.

### Flow cytometry

Cells were pelleted and washed with 250uL wash buffer (HBSS + 1% BSA, filtered with a 50mL Steriflip unit (Millipore, Burlington, MA)) prior to a 30m incubation in Fc Receptor Blocker (Innovex Biosciences, Richmond, CA) on ice in the dark. Cells were then washed twice before resuspension in wash buffer containing conjugated antibodies for flow cytometry at their recommended concentrations (Becton Dickinson, Franklin Lakes, NJ), and incubation for 30m on ice in the dark. Cells were then washed three times prior to flow cytometry. Doublets were called based on gates drawn for FSC-A and FSC-W, and dead cells were counted based on gates drawn in FSC-A and SSC-A. Gates for fluorescent markers were selected based on the background fluorescence of unstained controls, such that <5% of nonspecific cells would be counted as positively stained. All flow cytometry or FACS was performed on the Aria IIu in the Center for Advanced Technologies at UC San Francisco.

### MethoCult methylcellulose colony forming assay

Cells were counted with the TC20 automatic cell counter (Bio-Rad) with Trypan blue prior to plating in triplicate (as technical replicates) and incubation for 2 weeks at 37°C with 5% CO_2_. Colonies were then enumerated using MethoCult H4034 Optimum (Stemcell Technologies, Vancouver, CAN) according to the manufacturer’s instructions. Results shown are representative results from two independent experiments.

### Cell viability assays

For sorted drug sensitivity assays, cells were sorted with gates as previously described, and treated with drug. After 72h, dead cells were counted based on gates in FSC-A and SSC-A. For effector viability assays, cells were plated as technical replicates at 1e5 cells/mL with a WellMate (Thermo Fisher Scientific) and treated with drug or effector for 72h unless otherwise noted. Effectors were added with the Echo 525 (Labcyte, San Jose, CA). For readout, plates were allowed to cool to room temperature before combining well-mixed cells and cell media 1:1 with room temperature CellTitre-Glo 2.0 (Promega, Madison, WI) in opaque white tissue culture plates (Corning, Corning, NY). Reactions were allowed to proceed according to the manufacturer’s protocol, and luminescence was read out with the Biotek H4 plate reader (BioTek, Winooski, VT) in the Center for Advanced Technology at UC San Francisco. Results shown are representative results from three independent experiments.

### External marker tracking by flow cytometry

We used FACS to segregate equal numbers of the CD38^hi^ and CD38^lo^ subpopulations of Tet2^WT^ and Tet2^KO^ cells and seeded them separately in a round-bottom 96-well plate (CELLSTAR, Dallas, TX). Wells were treated with 0, 1, or 4uM AraC in triplicate (technical replicates), thus providing 9 initial conditions per sorted population per cell line. Samples were then re-profiled by flow cytometry for CD34 and CD38 every 3 days, unless otherwise noted. To account for day-to-day variations in Aria IIu laser power or other settings, the median intensity of unsorted, unstained controls of the appropriate cell line for each day was subtracted from raw fluorescence intensity of each sample. CD38^hi^ and CD38^lo^ subpopulations for analysis and model-fitting were called based on a threshold set midway between the gates used for sorting. Cell-state compositions of the samples served as input to the model to fit parameters for the cell-state transition matrix in both cell lines for both treated and untreated conditions.

### Data availability

RNA sequencing and DNA methylation data that support the findings of this study will be deposited in GEO and released upon publication. Code will be made available on GitHub (https://github.com/lmorinishi) upon publication. All other data are available from the corresponding authors upon request.

## Tables with Titles and Legends

**Supplemental Table 1.** Top Enrichr results from “ARCHS4 Tissues” for genes strongly correlated with Tet2 expression (Pearson correlation > 0.45) in the LAML study.

| Term | p-value | Adj p-value | Combined Score |
| --- | --- | --- | --- |
| GRANULOCYTE | 5.80E-35 | 6.27E-33 | 351.221812 |
| NEUTROPHIL | 2.87E-26 | 1.55E-24 | 229.673308 |
| DENDRITIC CELL | 4.19E-23 | 1.51E-21 | 189.887693 |
| PERIPHERAL BLOOD | 1.37E-21 | 3.69E-20 | 171.757628 |
| ALVEOLAR MACROPHAGE | 2.09E-19 | 4.52E-18 | 146.67615 |
| BLOOD DENDRITIC CELLS | 6.88E-13 | 1.24E-11 | 80.1008484 |
| MACROPHAGE | 3.58E-11 | 5.53E-10 | 64.8236466 |
| CORD BLOOD | 1.28E-07 | 1.73E-06 | 36.6594651 |
| SPLEEN (BULK TISSUE) | 6.91E-06 | 8.29E-05 | 24.8371736 |
| CD19+ B CELLS | 9.91E-04 | 0.01070011 | 12.1748187 |
| PLASMACYTOID DENDRITIC CELL | 0.00389624 | 0.03825395 | 9.15439163 |

**Supplemental Table 2.** Top Enrichr results from “ARCHS4 Tissues” for genes strongly correlated with Tet2 expression (Pearson correlation > 0 45) in the OHSU study

| Term | p-value | Adj p-value | Combined Score |
| --- | --- | --- | --- |
| NEUTROPHIL | 8.78E-22 | 9.49E-20 | 269.662309 |
| GRANULOCYTE | 1.18E-20 | 6.39E-19 | 248.485518 |
| PERIPHERAL BLOOD | 1.36E-13 | 4.90E-12 | 130.08582 |
| DENDRITIC CELL | 5.28E-11 | 1.42E-09 | 93.5226046 |
| MACROPHAGE | 5.84E-08 | 1.26E-06 | 56.0686849 |
| ALVEOLAR MACROPHAGE | 2.84E-07 | 5.12E-06 | 48.5367869 |
| BLOOD DENDRITIC CELLS | 1.29E-06 | 1.99E-05 | 41.6825623 |
| CORD BLOOD | 1.29E-06 | 1.74E-05 | 41.6825623 |
| SPLEEN (BULK TISSUE) | 0.00241362 | 0.02896349 | 13.2313735 |

**Supplemental Table 3.**
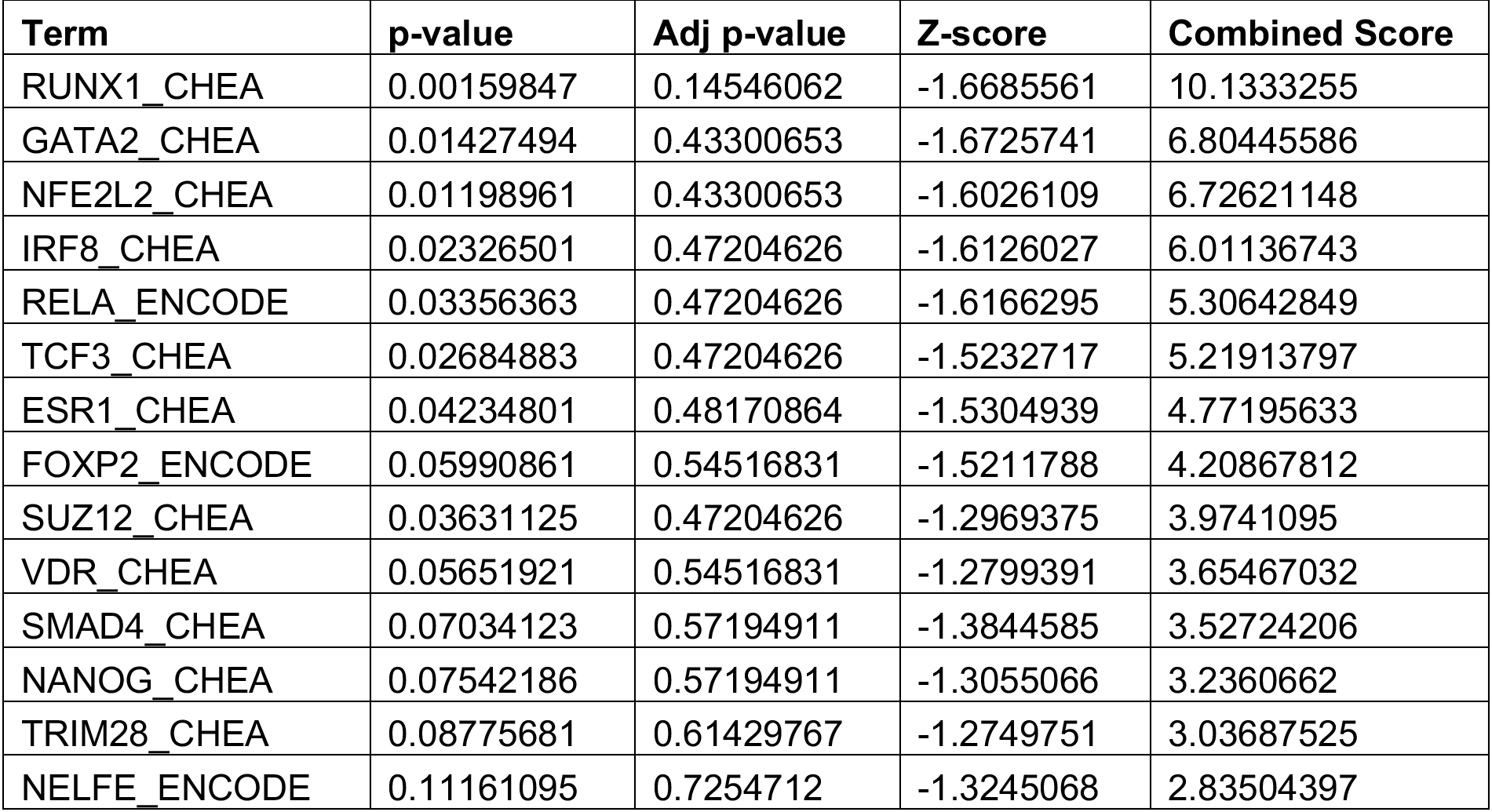
Top 15 Enrichr results from “ChEA 2016” for significantly differentially expressed genes in Tet2^KO^ AML cells compared to Tet2^WT^.

**Supplemental Table 4.**
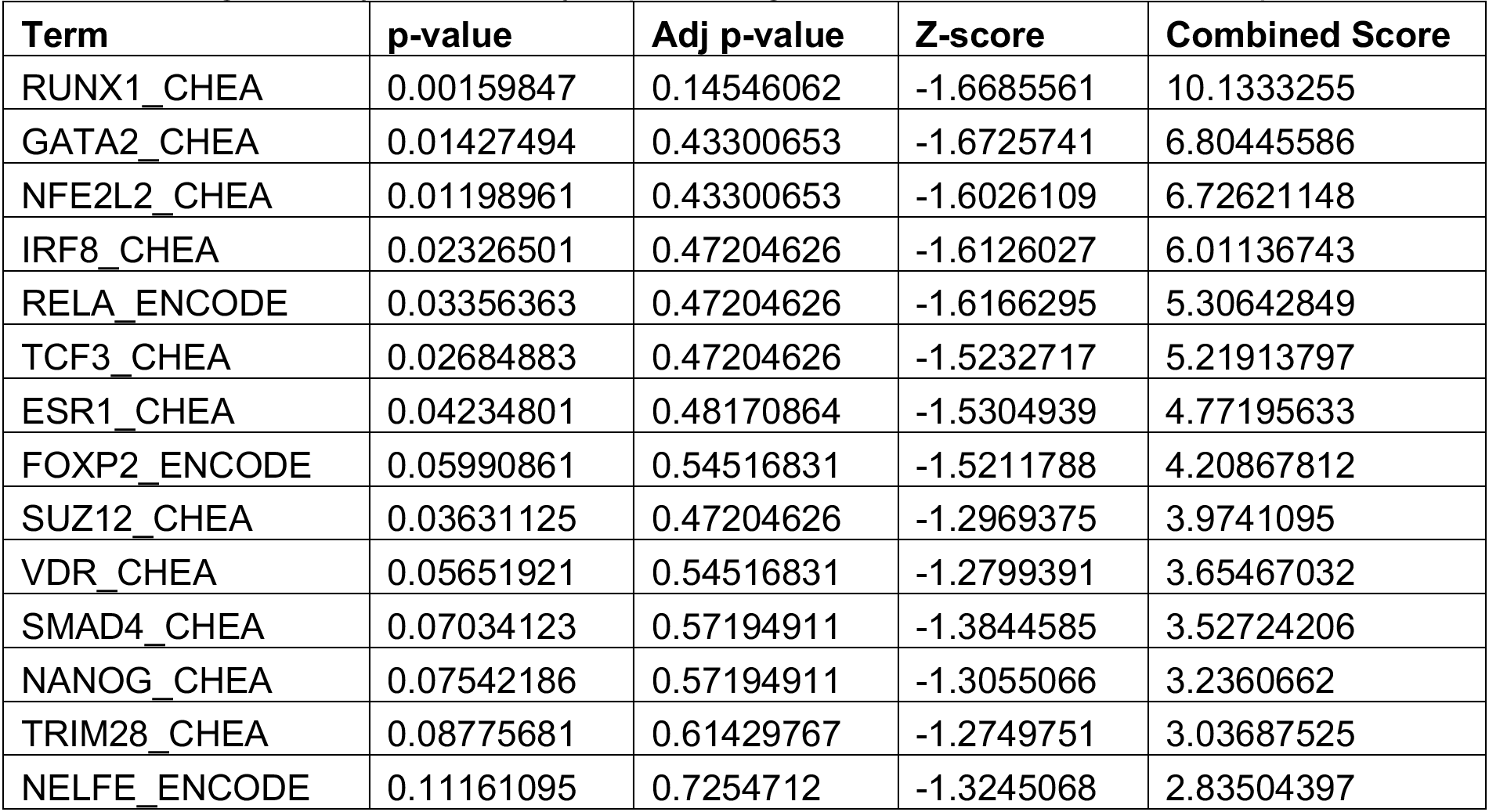
Top 15 Enrichr results from “ENCODE and ChEA Consensus TFs from ChIP-X” for significantly differentially expressed genes in Tet2^KO^ AML cells compared to Tet2^WT^.

**Supplemental Table 5.** Panel of effectors used to modulate chemosensitivity and halt the CD38^hi^ to CD38^lo^ transition.

| **Name** | **Category** | **Max concentration** |
| --- | --- | --- |
| DMSO | vehicle | -- |
| 2-deoxy-D-glucose | glycolysis | 2 uM |
| CCCP | oxphos | 50 nM |
| Disulfiram | redox | 1 nM |
| Etacrynic Acid | redox | 100 nM |
| IFNg | signaling | 50 ng/mL |
| Torin | AA | 1 nM |

## Supplemental Figure Titles and Legends

**Supplemental Figure 1.**
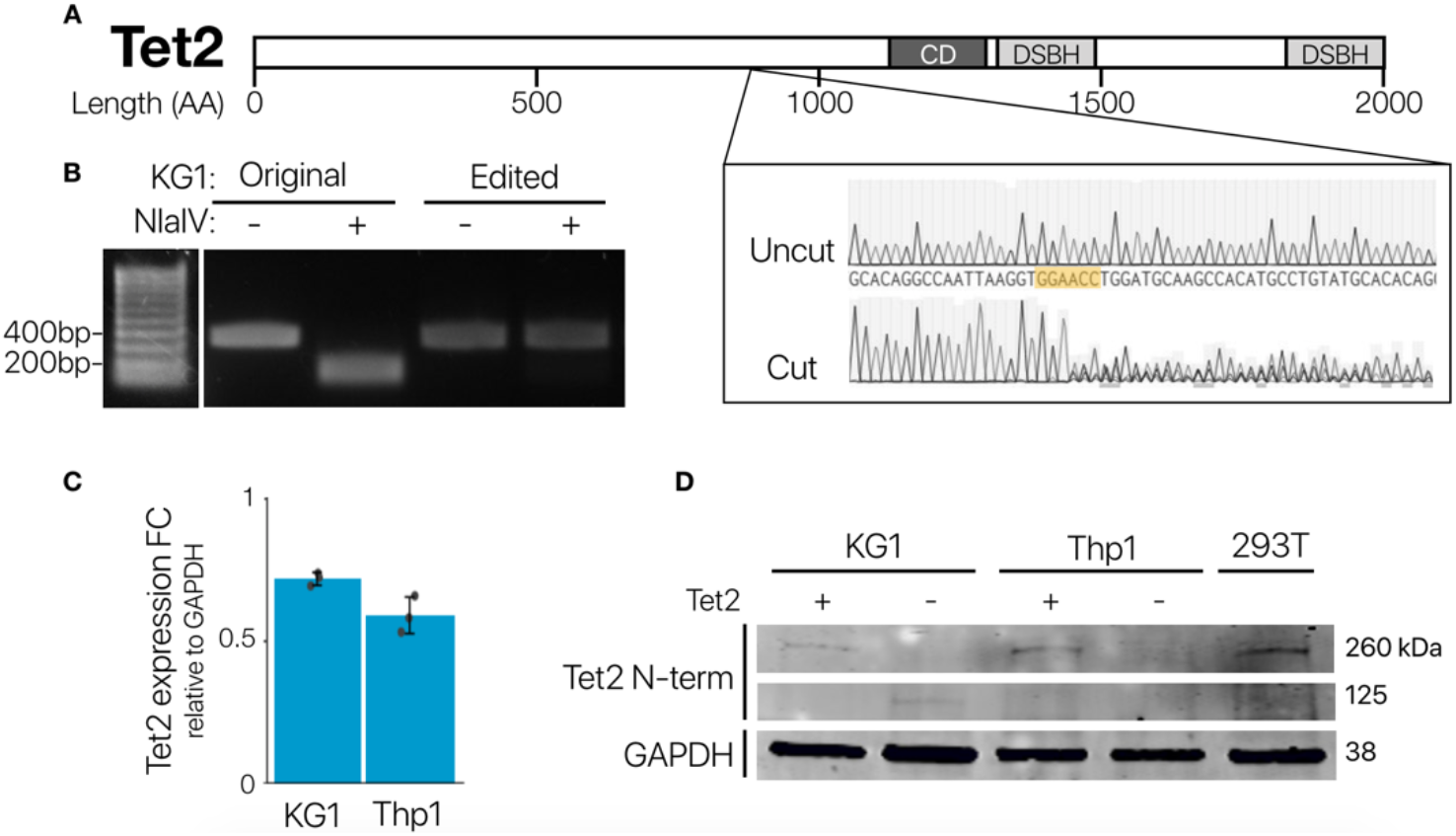
Confirming loss of Tet2 expression. (A) An outline of the gene Tet2 showing sequence the final gRNA targeted, and a bulk Sanger sequencing trace of the locus before and after plasmid nucleofection. (B) Upon editing, a NlaIV cut site is removed. PCR amplicons of a 400bp region around the target locus were run on an agarose gel with or without NlaIV. (C) Tet2 expression fold-change, as measured by RT-qPCR in Tet2^KO^ pool relative to the parental cell line, shows decrease in expression relative to GAPDH fold-change. (D) Western blot of both cell lines before and after Tet2 knockout with Tet2 N-terminus targeting antibody. 293T is a positive control. Tet2 band is lost at 260kDa.

**Supplemental Figure 2.**
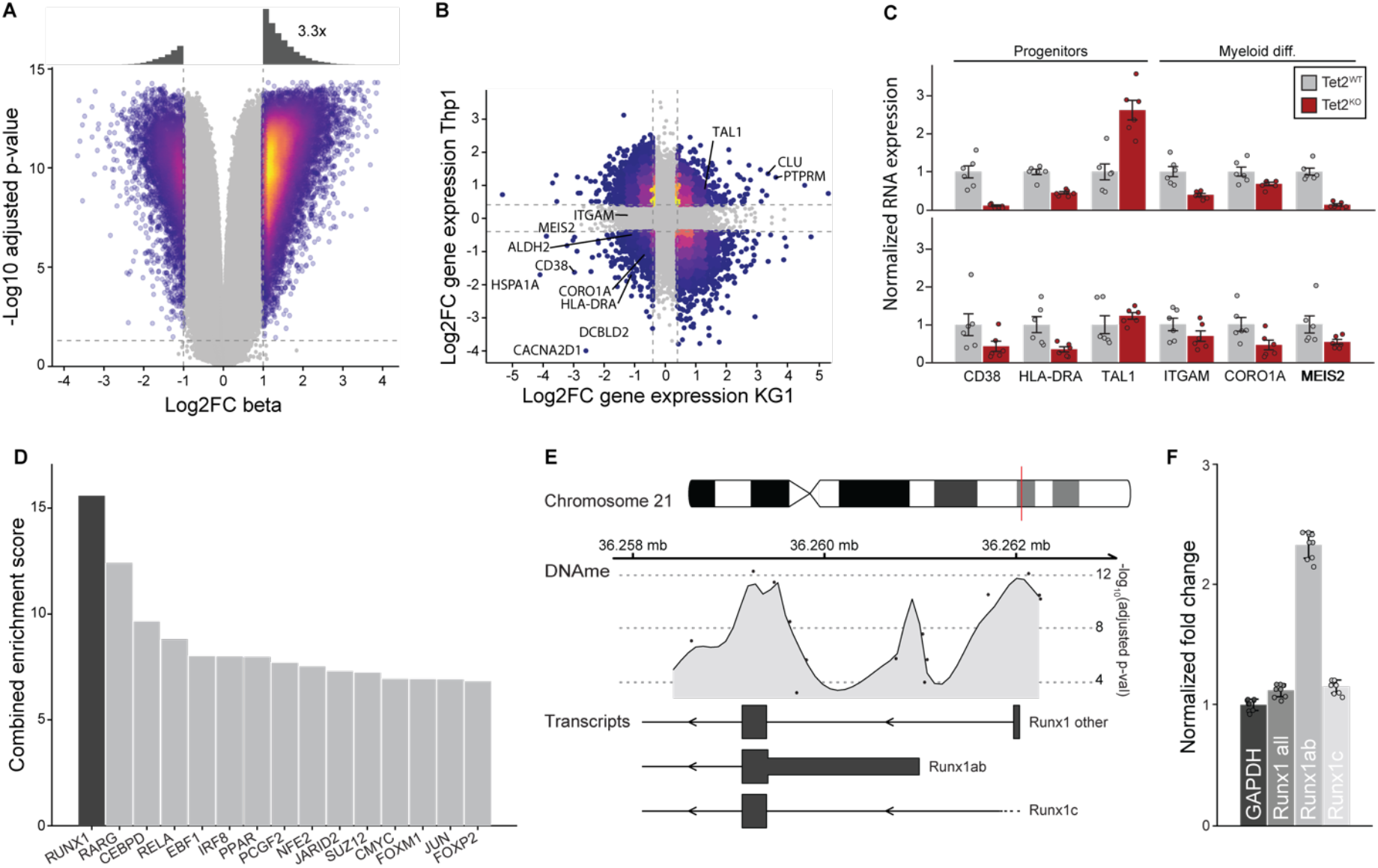
Changes in DNA methylation and RNA expression in Tet2^KO^ cells. (A) Representative volcano plot of log_2_fold-change in DNA methylation beta values after Tet2 loss (shown for KG1). Each point is one CpG. As expected, Tet2^KO^ cells are more hypermethylated at CpGs across the genome. (B) Log_2_fold-change of RNA expression counts from RNAseq after Tet2 loss for KG1 (x-axis) and Thp1 (y-axis). Each point represents a gene, and some similarly differentially expressed genes are labeled. Dotted lines denote a threshold of 1.5X change. (C) Genes that modulate differentiation and stemness are differentially expressed as measured by RNAseq after Tet2^KO^ in both KG1 and Thp1 (Supplemental Figure 2C, mean ± s.e., n=6). Tet2^KO^ cell lines show decreased expression of myeloid commitment markers (ITGAM, CORO1A) and increased expression of markers associated with stemness (CD38^-^, HLA-DRA^-^, TAL1) compared to Tet2^WT^. (D) Genes that are differentially expressed after Tet2 loss are enriched for targets of Runx1. Results from the “ChEA 2016” enrichment analysis of significantly differentially expressed genes after Tet2^KO^, showing transcription factors in order of “Combined Enrichment Score”. (E) Analysis of DNA methylation data found that the most significant DMR in Tet2^KO^ cells was the proximal promoter of Runx1. An ideogram representing the chromosomal location of Runx1 (top), the significance of differential methylation in the region (middle), and relevant transcripts (bottom) are shown. (F) Runx1a isoforms are differentially expressed in Tet2^KO^ cells. Fold-change of Runx1 isoforms in KG1 Tet2^KO^ cells is shown, normalized to GAPDH fold-change, calculated from a RTqPCR experiment (mean ± s.e., n=8).

**Supplemental Figure 3.**
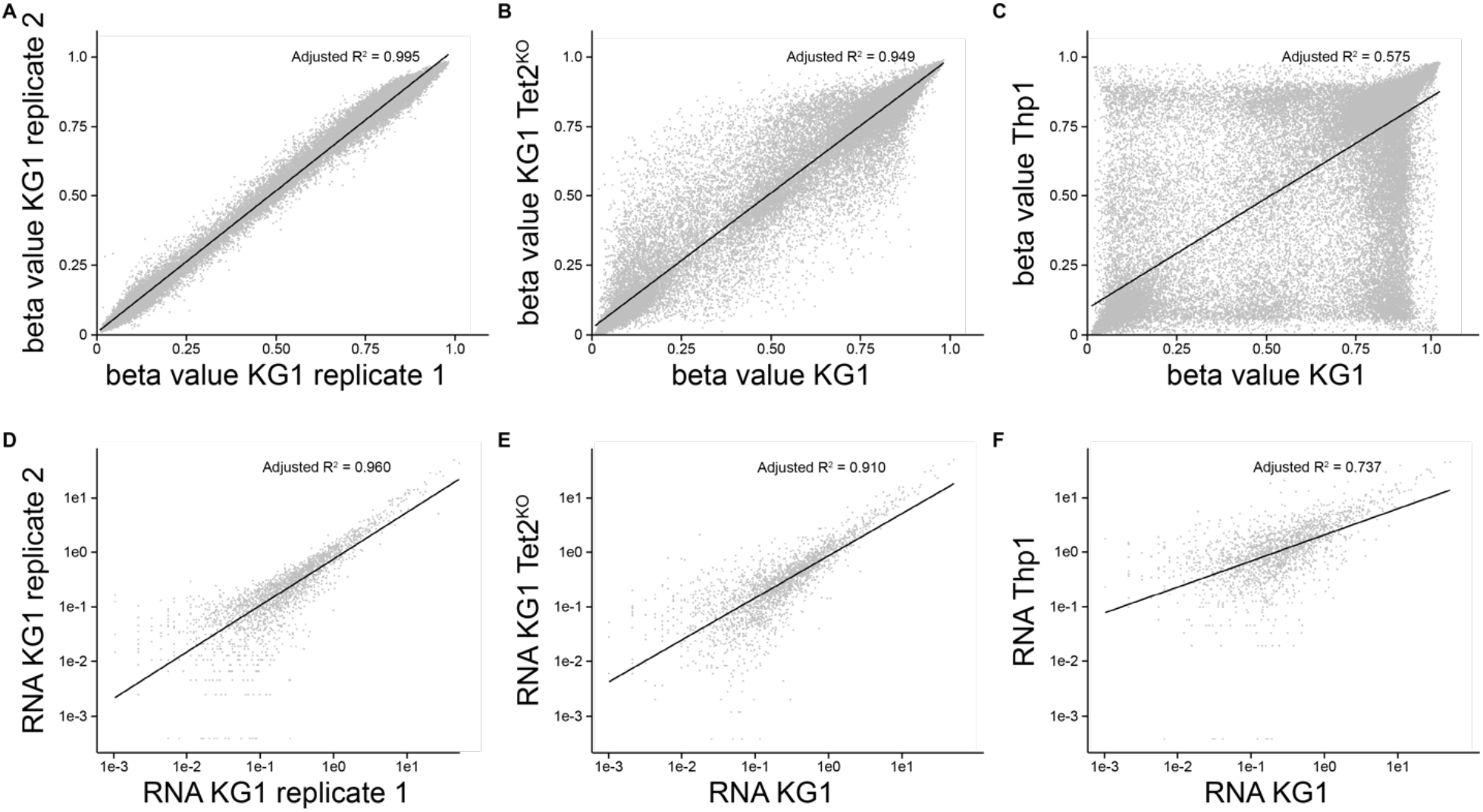
Correlation of DNA methylation and RNA expression across replicates, treatment conditions, and cell lines. Representative DNA methylation (A-C) and RNA expression (D-F) correlation between (A,D) technical replicates, (B,E) Tet2^WT^ and Tet2^KO^, and (C,F) different untreated cell lines.

**Supplemental Figure 4.**
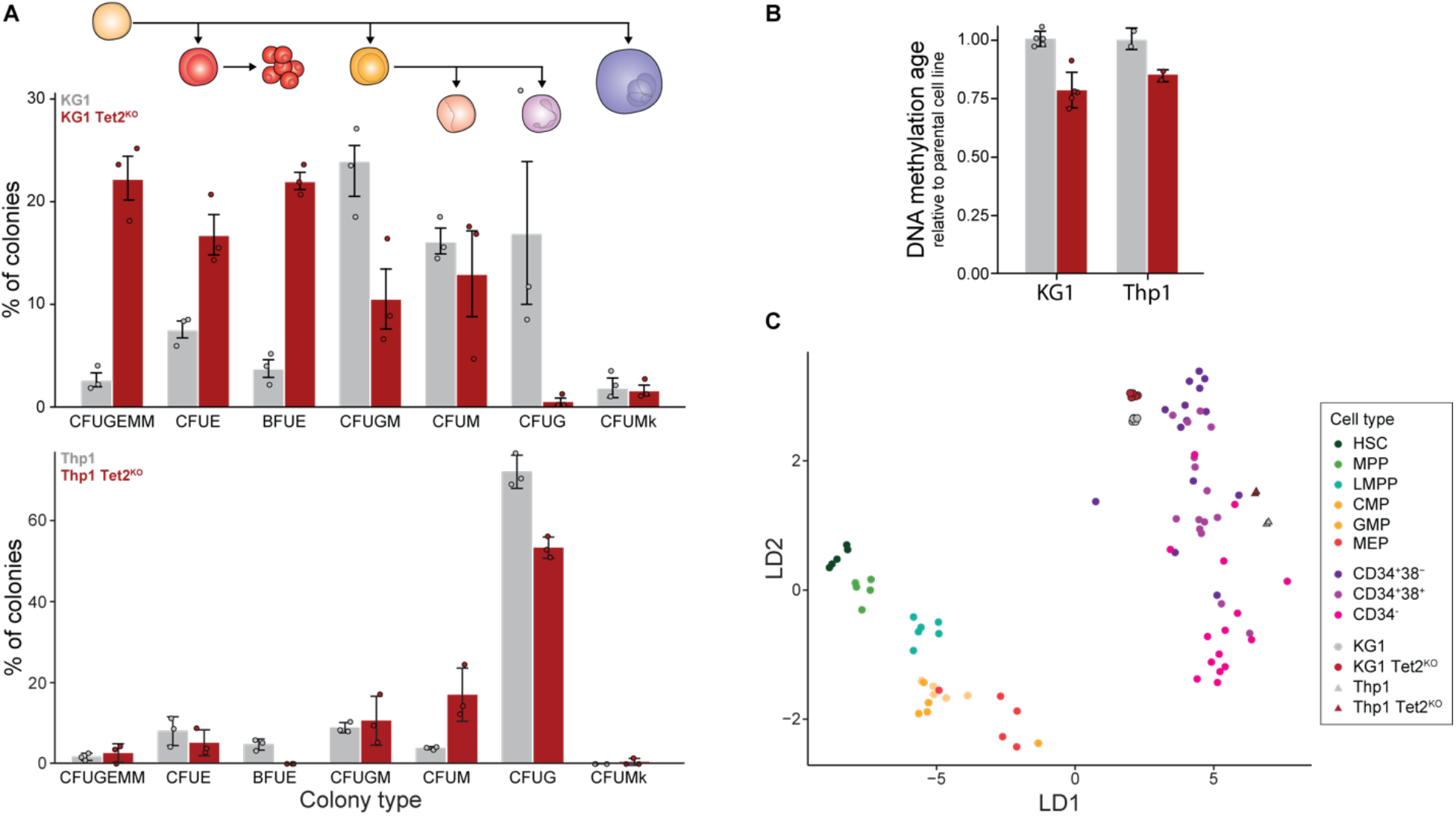
Tet2^KO^ myeloblasts are more stem-like than their Tet2^WT^ counterparts. (A) KG1 and Thp1 (Supplemental Figure 4A) Tet2^KO^ cell lines produce a higher percentage of colonies associated with oligopotent progenitor cells (CFUGEMM) compared to Tet2^WT^ cells (mean ± s.e., n=3) in methylcellulose colony forming assays. (B) DNA methylation age as measured by the Horvath calculator for KG1 and Thp1 cells for Tet2^WT^ (gray) and Tet2^KO^ (red) (mean ± s.e., n=3). (C) Low-dimensional representation of DNA methylation profiles after using LDA to find the space that best separates HSCs, CD34^+^38^-^, and CD34^+^ cells in a reference dataset^20^. Each point is one sample, and the color corresponds to the cell type of the sample. The two LD axes are readily interpretable: LD1 separates normal hematopoietic cells from leukemic and increasing LD2 moves up the known hematopoietic hierarchies. KG1 and Thp1 Tet2^KO^ cells have greater LD2 values than their corresponding Tet2^WT^ counterparts.

**Supplemental Figure 5.**
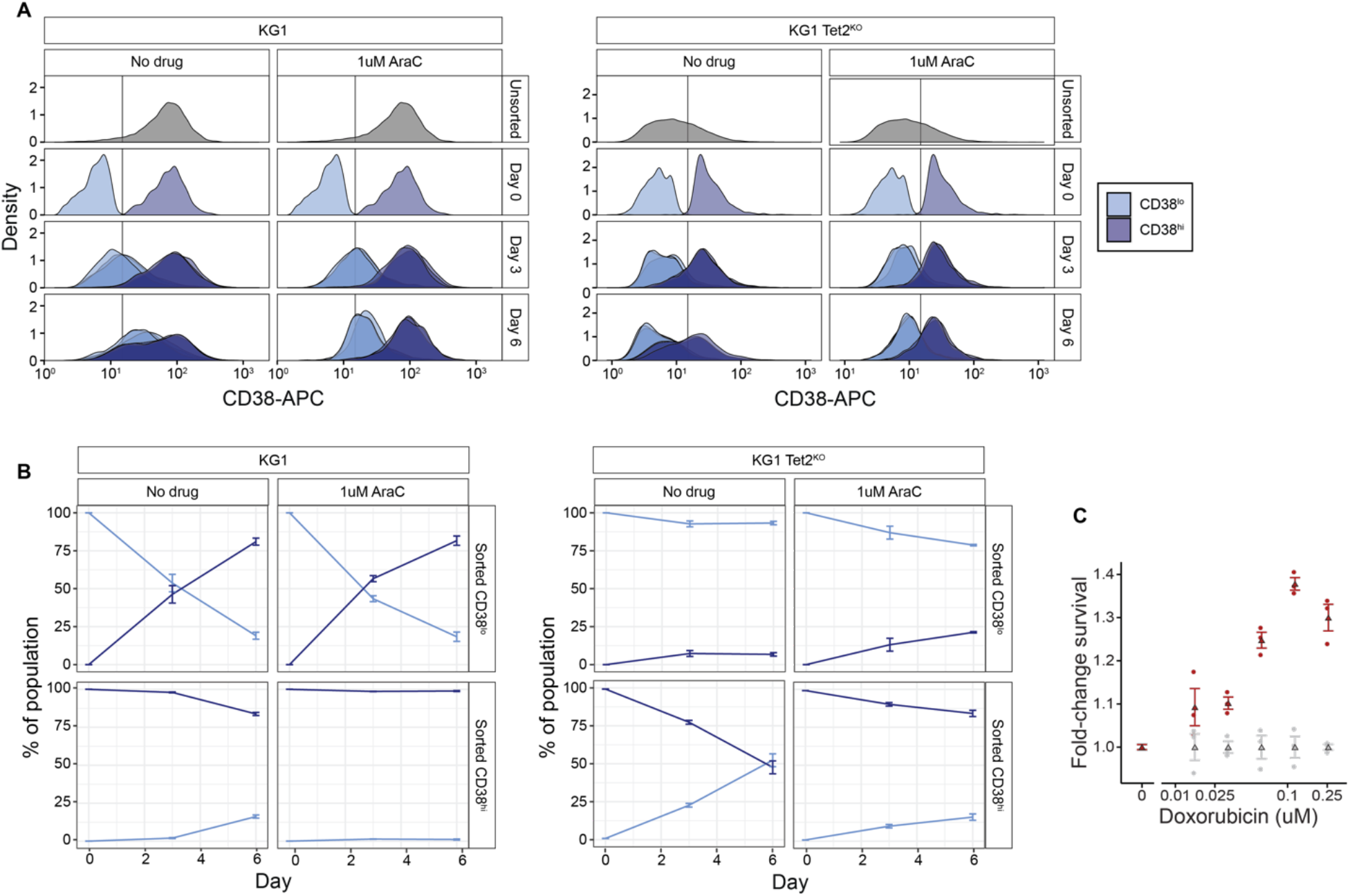
Tet2 status and drug treatment alter the dynamics of CD38 surface marker expression. (A) On day 0, cells from both KG1 and KG1 Tet2^KO^ cell lines (gray) were sorted based on CD38 expression (light blue: low, dark blue: high), and plated at equal densities. Every 3 days, cells were reflowed to quantify CD38 expression. Shown are the densities of the flowed populations, colored by the original sorted population (n=3). (B) Quantification of KG1 and KG1 Tet2^KO^ cells, respectively, called as CD38 high or low over time, based on the threshold (black vertical line in A, mean ± s.e., n=3). (C) The viability of cells treated with 72h of varying concentrations of Doxorubicin were assessed by CellTitreGlo. Shown is fold-change viability relative to Tet2^WT^ cells as a function of chemotherapy concentration (mean ± s.e., n=3).

**Supplemental Figure 6.**
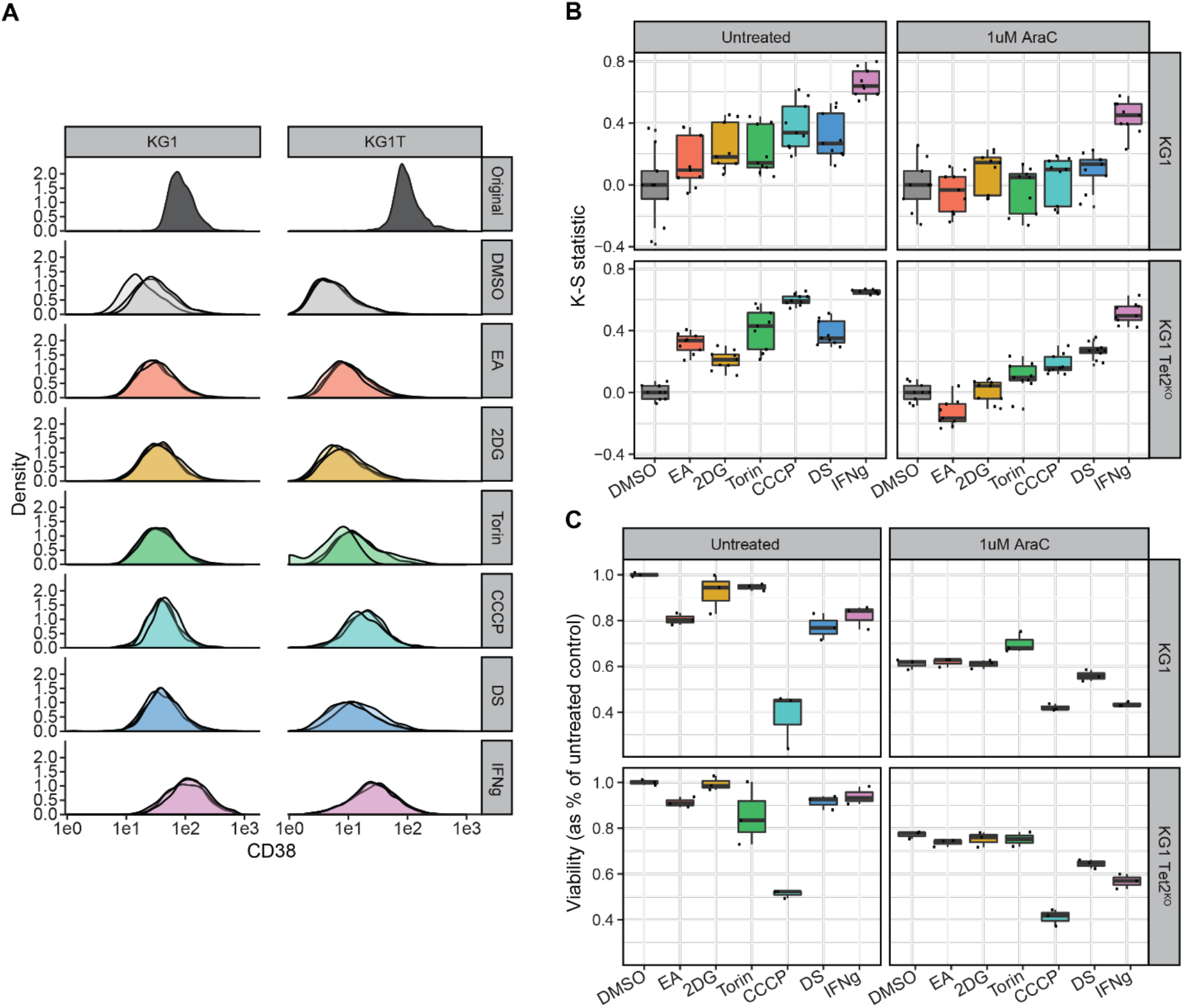
Testing a panel of effectors for cell state transition and chemosensitivity modulation. (A) KG1 and KG1 Tet2^KO^ CD38^hi^ cells were isolated and plated in triplicate. The dark gray density shows the CD38^hi^ population reprofiled just after sorting. For 3 days, CD38^hi^ samples were treated with effectors, in the presence or absence of 1uM AraC treatment (untreated condition not shown). Samples were then reprofiled with flow cytometry to assess how CD38 surface marker expression had changed relative to DMSO control (light gray density, n=3). (B) The signed Kolmogorov-Smirnov statistic was calculated to quantify differences in CD38 surface marker expression in effector treatment relative to DMSO control (n=3). (C) KG1 and KG1 Tet2^KO^ cells were treated with effectors, in the presence or absence of 1uM AraC treatment. After 3 days, viability was measured. Data shown is viability normalized to DMSO control in the untreated condition for each cell line (n=3).

